# The rat medial prefrontal cortex exhibits flexible neural activity states during the performance of an odor span task

**DOI:** 10.1101/454611

**Authors:** E De Falco, L An, N Sun, AJ Roebuck, Q Greba, CC Lapish, JG Howland

**Affiliations:** Department of Psychology, Indiana University-Purdue University Indianapolis, IN, USA; Department of Physiology, University of Saskatchewan, Saskatoon, SK, Canada; Present address: Medical College of Acupuncture-Moxibustion and Rehabilitation, Guangzhou University of Chinese Medicine, 510006 Guangzhou, China

**Author notes:** Corresponding authors: Health Sciences Building 107 Wiggins Rd Saskatoon, SK, Canada S7N 5E5 (t) 402 North Blackford St Indianapolis, IN, USA 46202 (t). These authors contributed equally to this work (E De Falco, L An).

**Keywords:** interneuron, metastable dynamics, nonmatching-to-sample, pyramidal neuron, working memory

## Abstract

Medial prefrontal cortex (mPFC) activity is fundamental for working memory (WM), attention, and behavioral inhibition; however, a comprehensive understanding of the neural computations underlying these processes is still forthcoming. Towards this goal, neural recordings were obtained from the mPFC of awake, behaving rats performing an odor span task of WM capacity. Neural populations were observed to encode distinct task epochs and the transitions between epochs were accompanied by abrupt shifts in neural activity patterns. Putative pyramidal neuron activity increased significantly earlier in the delay for sessions where rats achieved higher spans. Furthermore, increased putative interneuron activity was only observed at the termination of the delay thus indicating that local processing in inhibitory networks was a unique feature to initiate foraging. During foraging, changes in neural activity patterns associated with the approach to a novel odor, but not familiar odors, were robust. Collectively, these data suggest that distinct mPFC activity states underlie the delay, foraging, and reward epochs of the odor span task. Transitions between these states enable successful performance in dynamic environments placing strong demands on the substrates of working memory.

## Introduction

Working memory (WM) refers to the ability to hold and manipulate information on-line during a delay for future use. Understanding the neurobiological bases of WM is critical as deficits in WM are a central cognitive symptom of brain disorders including schizophrenia (Barch and Smith 2008; Barch et al. 2012). WM can be broadly parsed into domains including goal maintenance, interference control, and capacity (Barch and Smith 2008; Barch et al. 2012). Each of these domains requires several cognitive functions, such as planning, executive control, resistance to distraction, task monitoring, and memory. Electrophysiological recordings performed in animals engaged in WM tasks have identified the types of computations that brain regions or networks contribute to these functions (Constantinidis et al. 2018; Lundqvist et al. 2018). Prior work has indicated that optimal working memory performance requires that ensembles of medial prefrontal cortex (mPFC) neurons track the various requirements of a task (e.g., epochs, rules) (Lapish et al. 2008, 2015; Durstewitz et al. 2010; Del Arco et al. 2017) which may facilitate performance monitoring and error detection in these networks (Hyman et al. 2017). Trial-specific information must also be maintained over a delay for optimal performance of WM tasks. The role of the mPFC in the maintenance of information across a delay has been extensively interrogated and two hypotheses have emerged. Initially, the identification of neurons that are persistently active during the delay suggested that mPFC may serve as a “buffer” to temporarily hold information (Goldman-Rakic 1996; Funahashi 2015). However, this view has evolved to suggest that mPFC is more important for directing cognitive resources/attention toward relevant neural circuits that likely play a defined role in the stimulus maintenance (Curtis and D’Esposito 2003; Postle 2006; Tsujimoto and Postle 2012; Lara and Wallis 2015). For example, we found that dynamic changes in theta power (mPFC and hippocampus) and increased mPFC unit phase locking to hippocampal theta during the delay period of a spatial WM task predicted subsequent performance during the test phase of the spatial WM task (Myroshnychenko et al. 2017). Therefore, the goal of this study was to further understand the contributions of the rodent mPFC to WM by measuring neural activity in a validated measure of WM, the odor span task.

Recently, the NIH-initiated Cognitive Neuroscience Treatment Research to Improve Cognition in Schizophrenia group nominated the odor span task (OST) for assessing WM capacity in rodents (Dudchenko et al. 2000, 2013; Young et al. 2007). The OST is an incremental nonmatching-to-sample task that closely resembles human working memory tasks that assess span, a rare characteristic for rodent WM tasks (Dudchenko et al. 2013). While performing the OST, rodents are required to dig for food rewards in scented bowls (Figure 1-B) and typically achieve spans of 8 to 15 odors in the task, although higher spans can be attained under some conditions. Performance of the OST depends on a distributed neural circuit including mPFC and dorsomedial striatum but not the hippocampus or posterior parietal cortex (Dudchenko et al., 2000; Davies et al. 2013, 2017; Scott et al. 2018). Span is also impaired by manipulations related to schizophrenia including treatment with NMDA receptor antagonists (Davies et al. 2013, 2017; Rushforth et al. 2011; Galizio et al. 2013) and maternal immune activation during pregnancy (Murray et al. 2017). However, to date, no studies have assessed the patterns of neural activity underlying OST performance.

**Figure 1:**
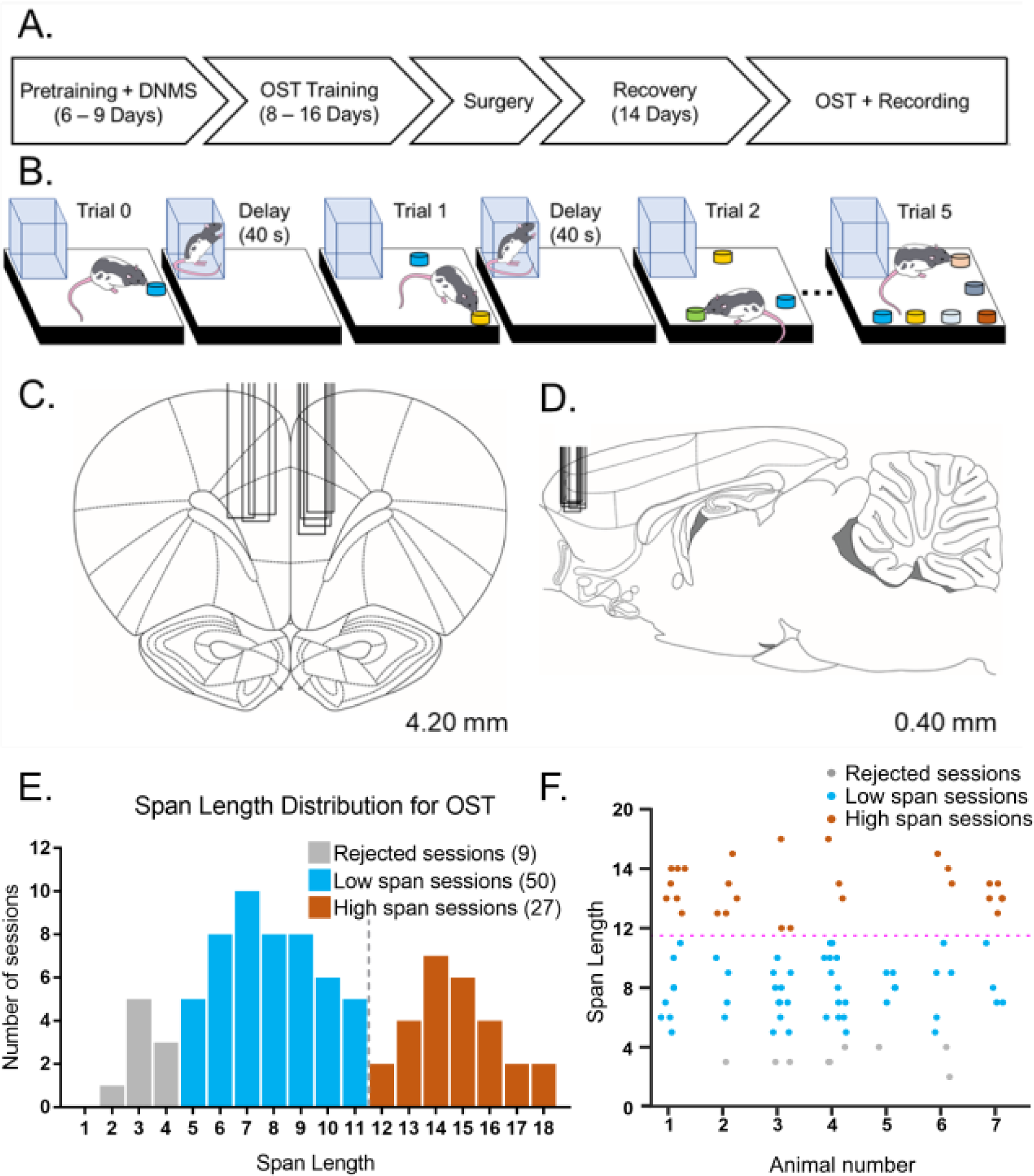
A- Timeline depicting experimental events. Pretraining and delayed non-match to sample training (DNMS) required 6-9 days of training. Training on the odor span task (OST) required 8-16 days of training. Following OST, animals underwent electrode implantation surgery and were allowed 14 days to recover. Following recovery, OST resumed, and electrophysiological recording occurred. B- OST consists of successive trials in which the animal must identify a novel odor and dig to receive a food reward. Different colors indicate different odors. With each successive trial, a new odor bowl (+) is added, while the previous odors (-) are rearranged pseudo-randomly. Between each trial the animal returns to a clear Plexiglas house for an intertrial delay period of about 40 s. OST continues until the animals fails to dig in the novel bowl. Span length is determined as the number trials successfully completed. C, D- Coronal and sagittal rat brain sections depicting the location of the recording sites. Probes were located in the prelimbic region of the mPFC. Box indicate the medial-lateral (panel C) and anterior-posterior (panel D) locations of the electrode arrays. E- Distribution of span lengths across the 86 recording sessions. The distribution is not unimodal (Calibrated Hartigan’s dip test, D(86) = 0.048, p=7.2×10^−3^). The local minimum between the two peaks (span = 11.5, black dotted line) was taken as threshold to classify the sessions into ‘Low span” (blue) and “High span” (red). Nine sessions with a span length smaller than 5 were excluded from the following analysis (grey). F- Span length for each session plotted by individual rats. Most rats (6/7) had both low and high span sessions.

Given the critical role of mPFC in the OST and the task’s similarity to human span tasks, we measured patterns of mPFC neural activity during performance of the OST in well-trained rats. In order to identify changes in neural activity required for optimal performance of the task, high span versus low spans sessions were analyzed during three task epochs: 1) the delay period, 2) the foraging period, and 3) the reward (or error) period. We predicted that delay-period activity of mPFC neurons recorded during the OST would predict span length in the OST. During foraging in the OST, rats approach familiar and novel bowls and sample the odors in exactly the same manner; however, when a novel odor is detected, they dig to retrieve the food reward. Thus, inhibition of digging is critical for performance of the task, particularly as the number of stimuli on the platform increases and the rats visit more familiar bowls during a bout of foraging. As proactive inhibition of motor responses is a recently described function of mPFC (prelimbic region) (Hardung et al. 2017), we anticipated to detect an inhibition signal during the foraging period. Finally, previous studies in decision-making tasks have shown error-related signals in mPFC neurons (Totah et al. 2009; Bissonette and Roesch 2015; Laubach et al. 2015), suggesting that this area is important for monitoring the outcome of actions during behavior. During the reward epoch of the OST, digging in the novel bowl enables retrieval of reward whereas the identical response (i.e., digging) in a familiar bowl is an error and results in the end of the session. Thus, by directly compared neural activity in these two types of trials, we were able to generate a pure error signal. To the best of our knowledge, no other task allows for a direct assessment of WM span or capacity in this manner. Thus, our results will information theories of mPFC function as the maximal working memory load (or capacity) is reached.

## Results

### Task normalization and the classification of neurons

A timeline of the experiment is illustrated in Figure 1-A. The OST, detailed in Figure 1-B, is designed to assess working memory capacity in rodents (Dudchenko et al. 2000). For each experimental session, the number of novel odors correctly identified (span length) is a measure of memory performance. In this experiment, 7 well-trained rats were implanted with mPFC electrodes and participated in a total of 86 recording sessions (see Materials and Methods and Table 1). The span length distribution across all recording sessions (Figure 1-E) was found to be bimodal; unimodality was rejected using a calibrated version of Hartigan’s dip test (Cheng and Hall 1998; Ardid et al. 2015) (Calibrated Hartigan’s dip test, D(86) = 0.048, p=7.2×10^−3^)The local minimum between the two peaks (span = 11.5) was then taken as threshold to separate ‘Low” and “High” span sessions. Nine sessions with span length smaller than five were excluded from all the following analysis because of the inadequate number of trials (Figure 1-F). Across the 77 recording sessions considered for analysis, following pre-processing and spike sorting, we identified 382 single neurons.

**Table 1.**
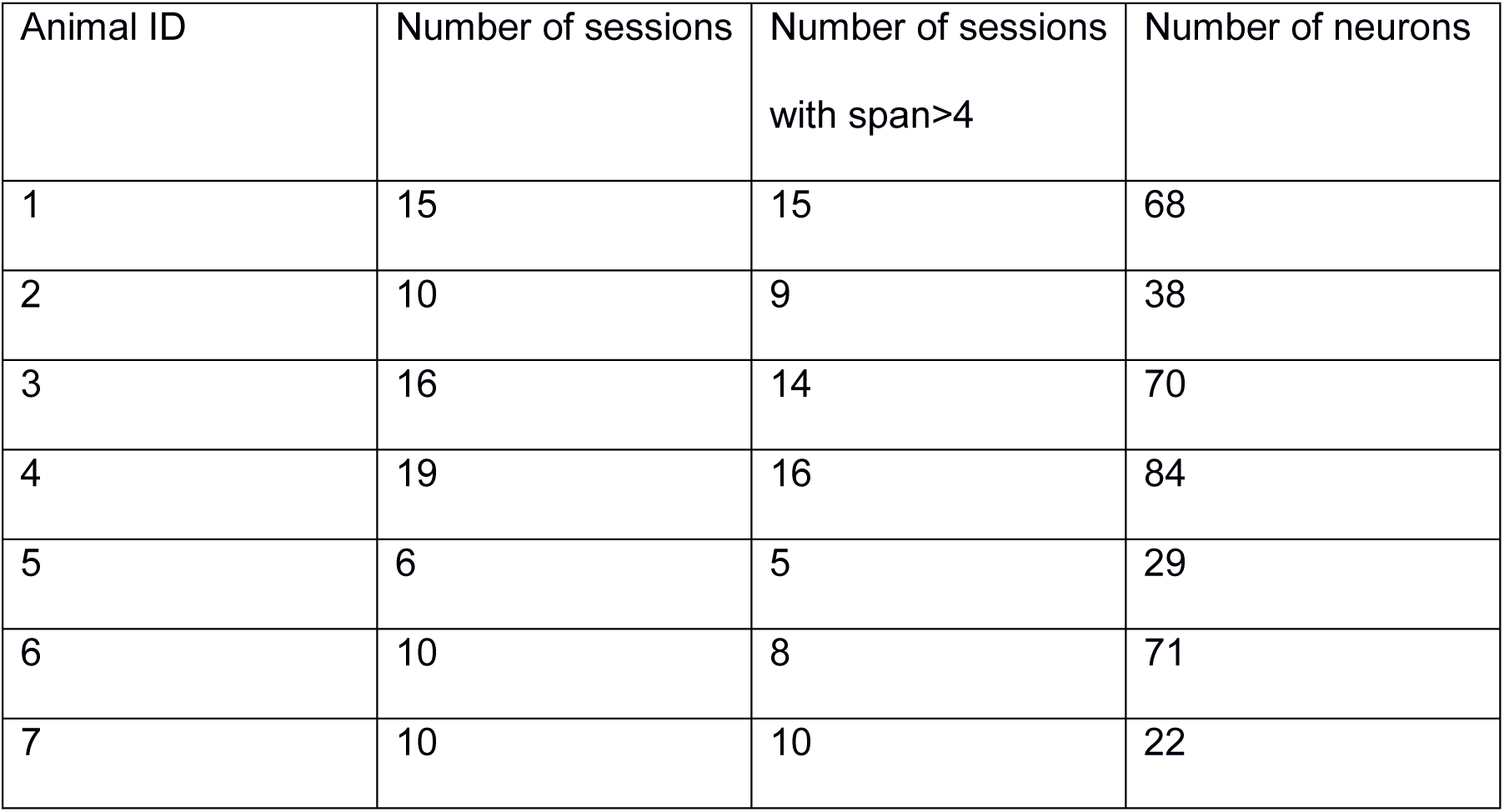
Number of recording sessions and number of neurons recorded for each animal.

| Animal ID | Number of sessions | Number of sessions<br>with span>4 | Number of neurons |
| --- | --- | --- | --- |
| 1 | 15 | 15 | 68 |
| 2 | 10 | 9 | 38 |
| 3 | 16 | 14 | 70 |
| 4 | 19 | 16 | 84 |
| 5 | 6 | 5 | 29 |
| 6 | 10 | 8 | 71 |
| 7 | 10 | 10 | 22 |

As each rat is permitted to forage at its own pace, the duration of foraging varies for each trail. To overcome this problem, we employed a targeted task-normalization of the firing rates. Figure 2-A2 shows a schematic of a single trial, where specific events were aligned to specific percentages of completion: the delay epoch covered from 0% to 70% of the trial, foraging epoch from 70% to 90%, and reward epoch (including the time to go back into the cage) from 90% to 100%. Through a time-normalized binning procedure (described in Materials and Methods) we obtained, for each neuron and each trial, a task-normalized firing rate trace spacing 100 bins, from the beginning to the completion of the task. For each neuron, the resulting firing rates were z-scored and then averaged across trials. The grand-average of normalized firing rates for the whole population of 382 neurons is shown in Figure 2-A1.

**Figure 2:**
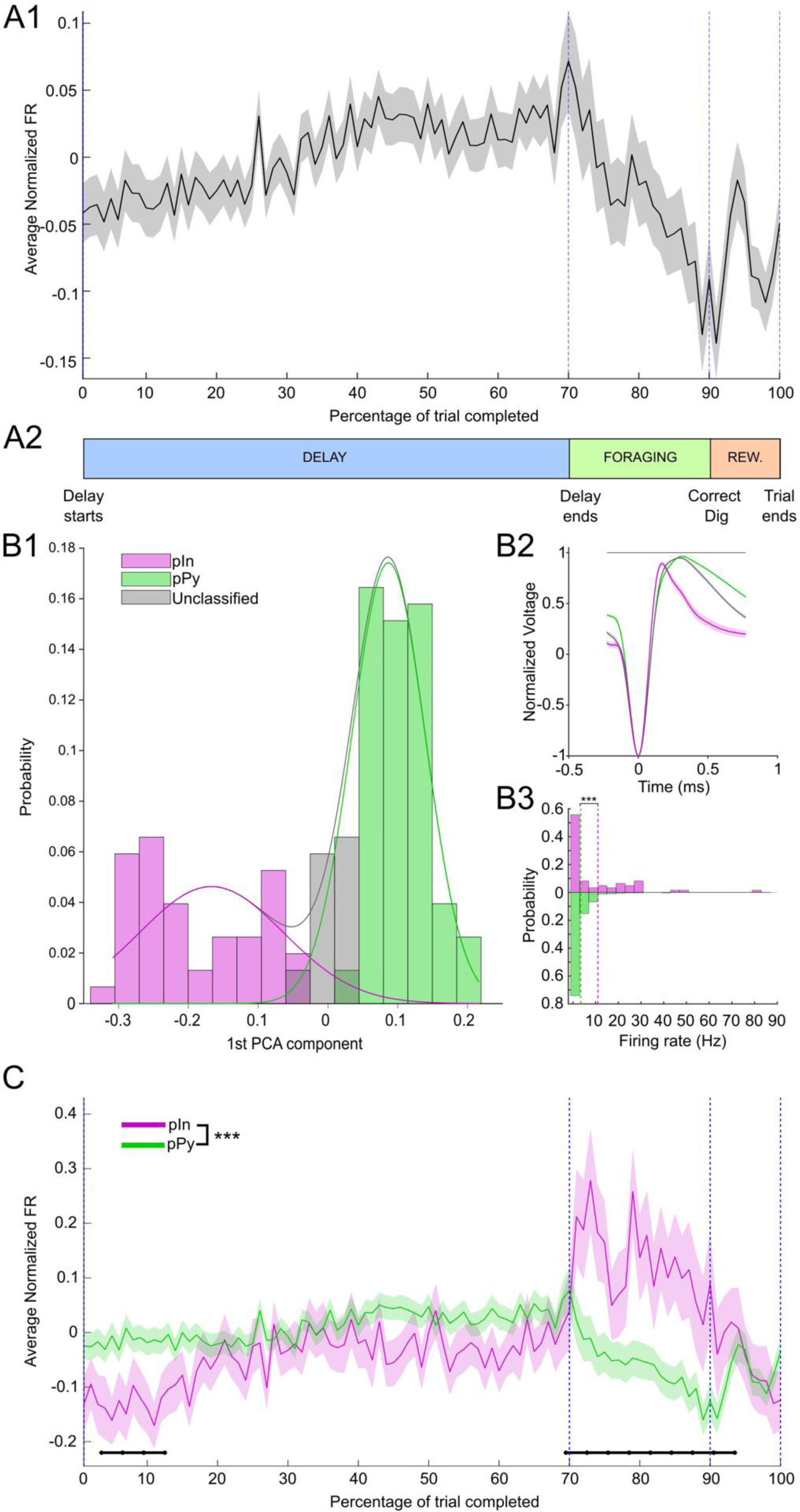
Task-normalized firing rates for pyramidal cells and interneurons. A1- Grand-average (±SEM) of task-normalized firing rate for 382 neurons recorded across 77 recording sessions. Firing rates were z-scored before averaging across neurons. A2- Timeline of a single trial, where the three main epochs of the task (Delay, Foraging, and Reward) were identified through the four behavioral timestamps: Delay starts; Delay ends; Correct dig; and End of trial. Specific percentages of completion were assigned to each task epoch to calculate the task-normalized firing rates (see Time Normalization in Materials and Methods). B1- Distribution of first PCA components (integrating two waveform features) for putative Interneurons (pIn), putative pyramidal cells (pPy) and unclassified neurons. The Gaussian fits used for the classification are shown as continuous lines on top pf the distribution. B2- Mean waveforms (±SEM) for the three classes of neurons detailed in panel B1. Unclassified neurons had a mean waveform closer to the pPy class and where subsequently labeled as pPys. B3- Distribution of mean firing rates for 61 pIns and 321 pPy. Firing rates were higher in the pIn population than in the pPy one (Kolmogorov-Smirnov test, D(321,61)=0.30, p=1.1×10^−4^). Vertical dotted lines mark the mean value of each distribution. C- Grand-average (±SEM) of task-normalized firing rate for pIns and pPys. The firing rates in the two classes were significantly different (2-way ANOVA, interaction cell class x time, *F*(99,38000) = 3.02, *p* = 8.4×10^−22^). Black horizontal lines mark groups of time bins with significant differences between pIns and pPy (FDR-corrected rank-sum, p<0.05).

Neurons were classified in putative Interneurons (pIn) and putative Pyramidal cells (pPy) according to their average waveform. The waveform features considered were integrated into the first component of a PCA and two Gaussian models were fit on the distribution of first components (Figure 2-B1) to separate narrow waveforms (corresponding to pIns) and broad waveforms (corresponding to pPy). Details about the classification procedure are described in Materials and Methods. Average waveforms and mean firing rate distribution for the classified cells are shown in panel B2 and B3 of Figure 2, respectively. At the end of the classification procedure 61 out of the 382 (16%) were classified as pIns, while the remaining 321 were classified as pPy. Firing rates across the population of pIns were significantly different from firing rates across the population of pPys (2-way ANOVA, interaction cell class x time, *F*(99,38000) = 3.02, *p* = 8.4×10^−22^), with significant differences (FDR-corrected rank-sum, p<0.05) during the first 13% of the task and during the whole foraging epoch (Figure 2-C). In particular, pIns exhibited a distinct pattern of activity, most prominently characterized by an increase in the average firing rate during the foraging and reward epochs compared to the delay epoch.

### Neural activity patterns robustly remapped between task epochs and predicted span

The most prominent neural activity patterns underlying the average firing rate profile were assessed via PCA. Figure 3-A shows the first three PCs identified. The task-normalized firing rates for the whole population of neurons were sorted according to their loadings on each of the first three PCs (Figure 3-B). From the sorted firing rates, distinct neural populations were observed whose firing rates changed together throughout the different task epochs. In particular, groups of foraging-active neurons and delay-active neurons emerged when sorting according to the first PC (left panel of Figure 3-B); neurons active during the first part of the delay epoch (early-delay) were separated from neurons active in the second part of the delay epoch (late delay) when sorting according to the second PC (middle panel of Figure 3-B); and neurons particularly active during the reward epoch were identified by the third PC (right panel of Figure 3-B). Remapping of neural activity signaled the transition between task epochs with a sharp and abrupt transition between delay and foraging epochs, and a smaller transition between foraging and reward epochs. Interestingly, pIns loaded more heavily than pPys on the first PC (Figure 3-C, left panel, Kolmogorov-Smirnov test: D(321,61)=0.24, p=4.9×10^−3^), indicating that interneurons mostly stayed active from the end of the delay and throughout the foraging epoch (in line with their average firing rate observed in Figure 2-C). No significant difference was observed for loadings on second and third PCs (Figure 3-C, middle and right panels, Kolmogorov-Smirnov test: D(321,61)=0.10, p=0.63 for PC2; D(321,61)=0.12, p=0.37).

**Figure 3:**
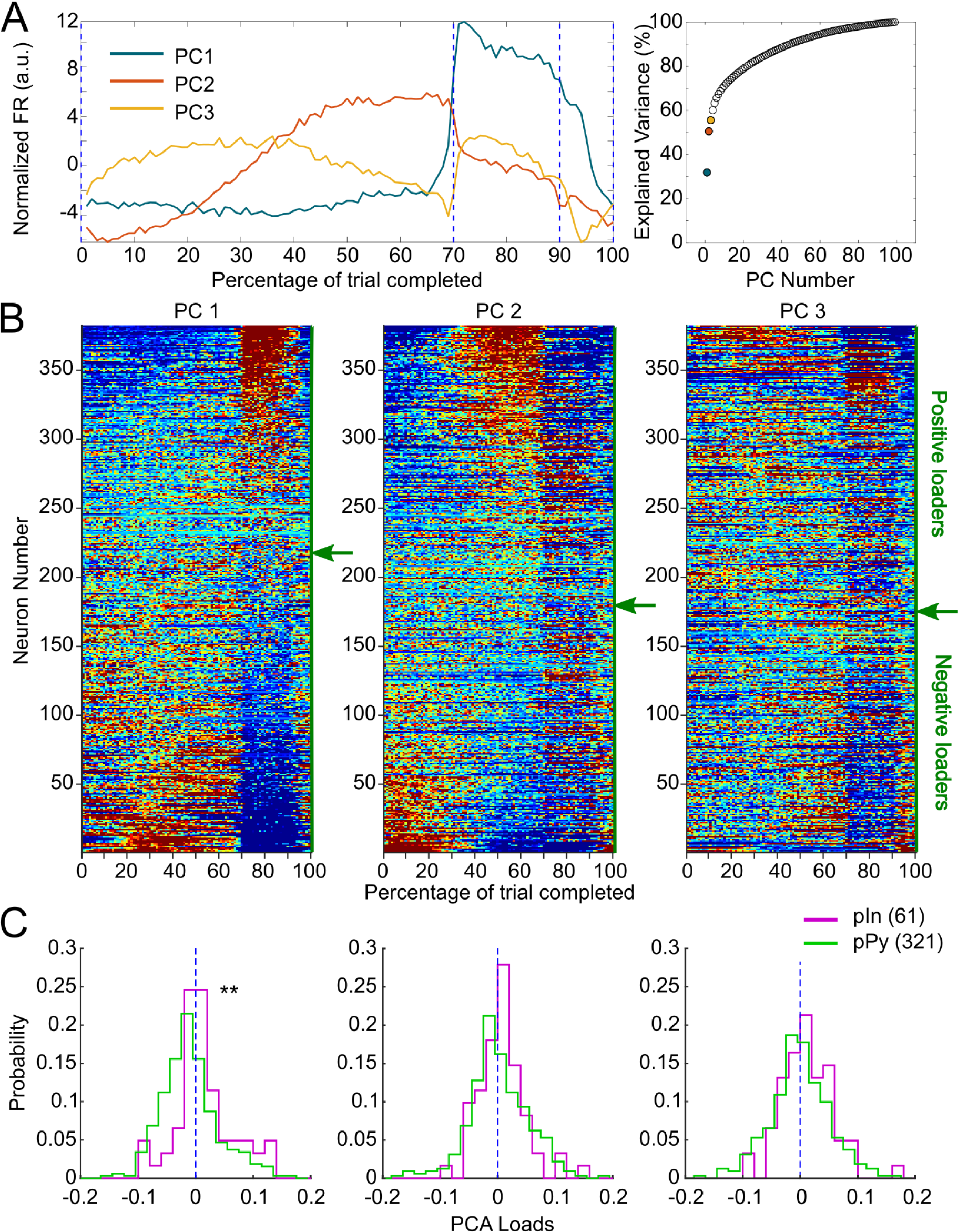
Identification of neural populations via PCA. A-First three principal components (PC). Projection of firing rates for the 382 neurons along the first three principal eigenvectors identified through PCA (left panel) and variance explained by each PC (right panel). The first three PCs together explained 56% of the original variance of the dataset. B-Task-normalized firing rates for the 382 neurons identified sorted according to their loadings on first, second and third PC (left, center, and right panel, respectively). Green arrows on the right side of each color-plot indicates the transition point between positive and negative loaders. C-Distributions of loadings on each PC separated for pIns and pPys. On the first PC pIns’ loadings were significantly higher than pPys’ ones (left panel, Kolmogorov-Smirnov test: D(321,61)=0.24, p=4.9×10^−3^), while no significant effect was found on the other two PCs (Kolmogorov-Smirnov test: D(321,61)=0.10, p=0.63 for PC2; D(321,61)=0.12, p=0.37).

We then examined the relationship between neural firing rates and odor span. Sessions were separated into high and low span (according to the threshold defined in Figure 1-C) and firing rates were compared for both pPys and pIns in the two span groups. The average firing rates (± SEM) separated by span group are reported in Figure 4-A and Figure 4-B for pPys and pIns, respectively. Firing rates of pPys recorded from low vs high span sessions were significantly different (2-way ANOVA, interaction span class x time, F(99, 31900) = 1.72, p=1.1×10^−5^), with bin-by-bin differences observed especially during the middle part of the delay epoch (FDR-correct rank-sum, p<0.05). No significant differences were observed in the firing rates of pIns when comparing low and high span sessions (2-way ANOVA, F(99, 5900) = 0.87, p=0.81).

**Figure 4:**
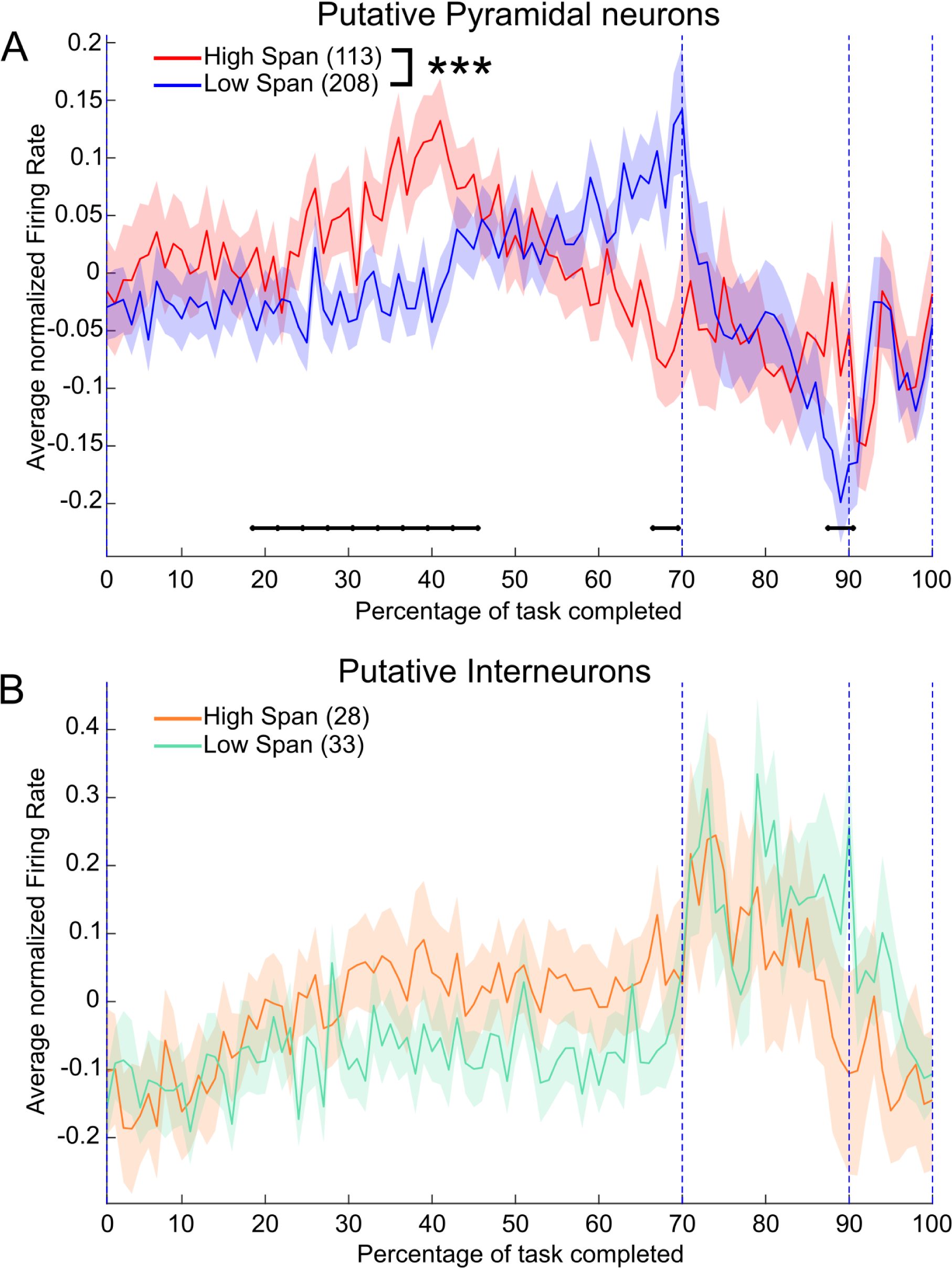
Activity of pyramidal neurons is predictive of span. A- Grand-average (±SEM) of task-normalized firing rate for 321 pPys, separated according to the session’s span (low and high span were defined according to the threshold identified in Figure 1-C). Firing rates in the two groups were significantly different (2-way ANOVA, interaction between span class and time bin, F(99, 31900) = 1.72, p=1.1×10^−5^). Black horizontal lines mark groups of time bins showing significant differences between low and high span groups (FDR-corrected rank-sum, p<0.05). B- Grand-average (±SEM) of task-normalized firing rate for 61 pIns, separated according to the session’s span. Firing rates in the two groups were not significantly different (2-way ANOVA, interaction between span class and time bin F(99, 5900) = 0.87, p=0.81).

Differences in neural activity patterns were observed in pPys across low and high spans. Since, as seen in Figure 3-B, different neurons exhibit different firing patterns throughout the execution of the task, we asked whether specific subsets of neurons were responsible for driving the difference between low and high span seen in Figure 4-A. We used a k-means clustering approach based on PCA-features (see Materials and Methods) to identify sub-classes of pPys responding in a similar manner to the different task epochs. The ideal number of clusters (k=4) was selected by means of the Akaike information criterion (Figure 5-A1). The 3-D feature space considered for clustering (loading on the first three PCs for each neuron) and the classification results are shown in Figure 5-A2. The average firing rate (±SEM) for each of the four classes identified is shown in Figure 5-A3.

**Figure 5:**
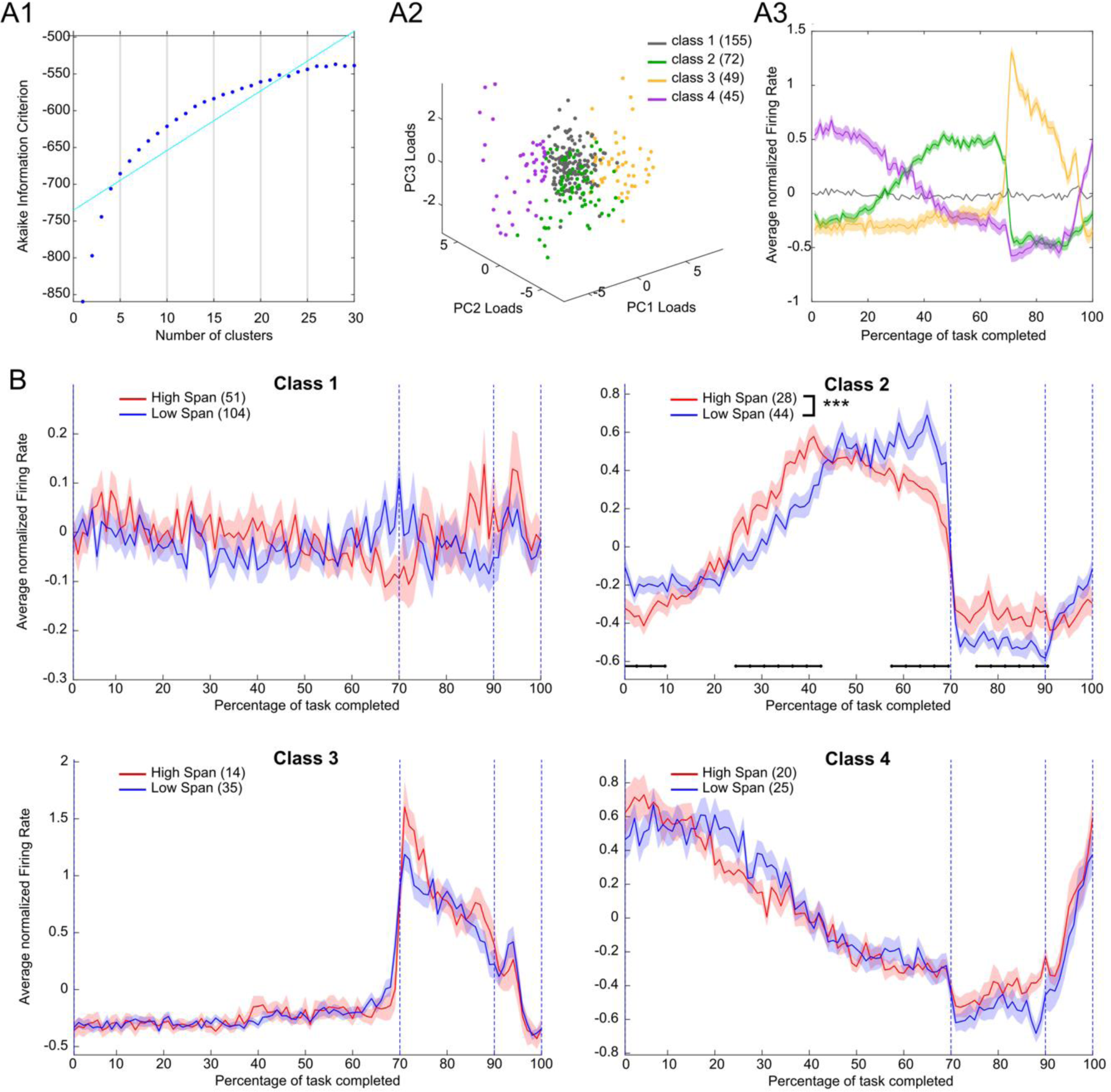
Identification of subpopulations of pyramidal neurons. A1-Akaike Information Criterion (AIC) for the PCA-features k-means clustering, calculated for different number of clusters (k). The selected number of clusters (k=4) was identified through a broken stick fit (cyan line). A2- Loadings on the first 3 PCs for the population of 321 pPys clustered. Different colors indicate the different classes assigned. A3- Average (±SEM) task-normalized firing rate for each of the classes identified. B- Grand-average (±SEM) of task-normalized firing rate for each class of pPys, separated according to the session’s span (low or high). Only firing rates in Class 2 were significantly different (2-way ANOVA, interaction span class x time, F(99, 7000) = 3.59, p=1.4×10^−29^). Black horizontal lines mark groups of time bins showing significant differences between low and high span groups (FDR-corrected rank-sum, p<0.05). No significant differences between firing rates in the low and high span sessions were observed in the remaining classes (2-way ANOVA, interaction span class x time: F(99, 15300) = 1.23, p = 0.06 for class1; F(99,4700) = 0.93, p = 0.67 for class 3; F(99,4300) = 1.18, p = 0.11 for class 4.

When looking at the relationship between span and firing rate for each of the subclasses identified (Figure 5-B), only Class 2 exhibited a significant difference between firing rates in the low vs the high span sessions (2-way ANOVA, interaction span class x time F(99, 7000) = 3.59, p=1.4×10^−29^), localized both during the delay and the foraging epochs (FDR-correct rank-sum, p<0.05). Neurons in this class were increasing their activity during the second half of the delay epoch. In the high span sessions, such increase started earlier and decreased more gradually towards the end of the delay compared to the low span sessions. No significant differences between firing rates in the low and high span sessions were observed in the remaining classes (2-way ANOVA, interaction span class x time: F(99, 15300) = 1.23, p = 0.06 for class1; F(99,4700) = 0.93, p = 0.67 for class 3; F(99,4300) = 1.18, p = 0.11 for class 4.

### Neural activity changes upon approach to novel, but not familiar, odors

During foraging, neural trajectories associated with the approach to familiar and novel odors were found to diverge right around the time of the approach (Figure 6-A). Trajectories were obtained through PCA on 188 neurons available (see Materials and Methods). Pronounced changes in neural activity were observed upon approach to a novel odor, while neural patterns associated with familiar odor approaches were weaker (Figure 6-B). The effect was statistically confirmed by comparing the distributions of absolute loadings on the first three PC’s for familiar vs novel approaches (Figure 6-C), Kolmogorov-Smirnov test: D(188,188)=0.22, p=2.0×10^−4^ for PC1; D(188,188)=0.19, p=2.5×10^−3^ for PC2; D(188,188)=0.15, p=2.7×10^−2^ for PC3). Note that both the pattern classification and the statistical comparison (Figure 6-B and 6-C) were performed by including only the neural activity up to 0.3 s after the approach in order to avoid contamination in the activity coming from the following events.

**Figure 6:**
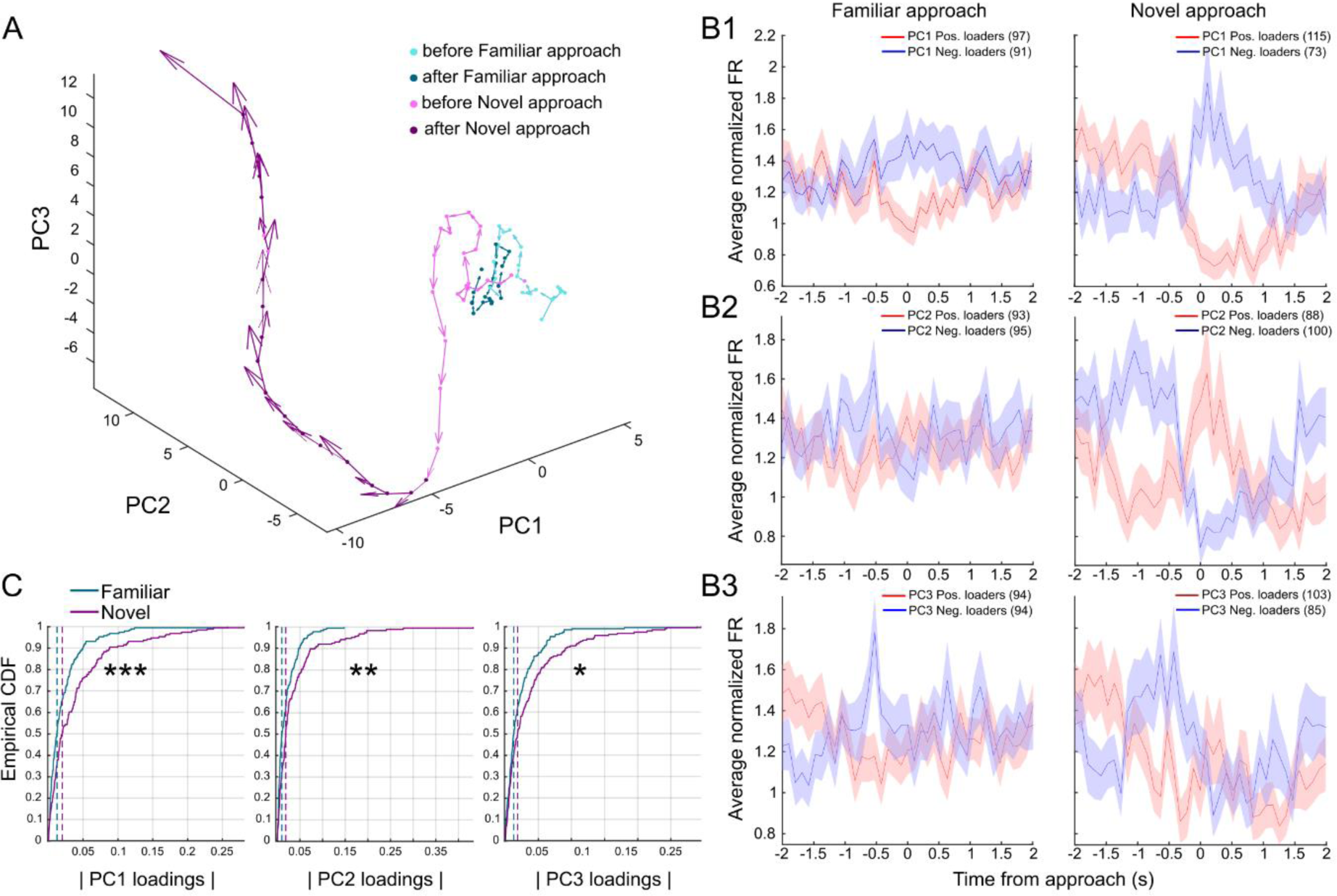
Distinct neural trajectories for familiar and novel odor approaches. A- Neural activity trajectories in the PC space for 188 pyramidal neurons around familiar and novel approaches (time interval −2 s to 2 s around each event, first 3 PC’s explaining 56% of variance). Arrows indicate module and direction of trajectories’ speed. B1-B2-B3- Average normalized firing rates (±SEM) for positive and negative PC loaders for familiar approaches (left panels) and novel approaches (right panels). Loadings were obtained considering a time interval from −2 s to 0.3 s around each event.C- Empirical cumulative distribution function (CDF) of absolute loadings on the first three PC’s for familiar and novel approaches (time interval −2 s to 0.3 s around each event, first 3 PC’s explaining 74% of variance). Absolute loading’ distributions in the two classes were different (Kolmogorov-Smirnov test: D(188,188)=0.22, p=2.0×10^−4^ for PC1; D(188,188)=0.19, p=2.5×10^−3^ for PC2; D(188,188)=0.15, p=2.7×10^−2^ for PC3).

The divergences in neural dynamics observed in different aspects of the task (i.e., related to performance during the delay epoch and to the approach to novel odors during foraging) led us to speculate whether differences in mPFC neural activity could be detected on trials where the animal dug in an incorrect bowl. A PCA of correct and incorrect trials was performed to address this issue (see Materials and Methods). Figure 7-A shows the neural trajectories for correct (shades of green) and incorrect (shades of red) trials in PC space. The neural trajectories provide a qualitative assessment of the most predominant population activity patterns in mPFC that are observed across the delay, foraging, and reward epochs. For correct trials the neural activity patterns follow a circular trajectory in this space, where the neural activity patterns at the end of the trial are similar to those observed in the start of the trial. However, on an incorrect trial, a sharp divergence in the trajectory’s direction was observed along PC3 at the beginning of the reward epoch, which corresponded to the activity of a group of neurons that started firing during this time on error trials only (top positive loaders on PC3, Figure 7-B). Firing rates for the top 30% of positive loaders on PC3 (30 pPys) were significantly different on correct vs incorrect trials (2-way ANOVA, interaction between kind of trial and time bin, F(99, 5800) = 1.59, p=2.0×10^−4^), with differences localized exclusively during the epoch following the dig (FDR-corrected rank-sum, p<0.05, Figure 7-C). Collectively, these data indicate that a unique signal emerges in mPFC immediately upon digging in an empty bowl that corresponds to a small subset of pyramidal neurons that start firing.

**Figure 7:**
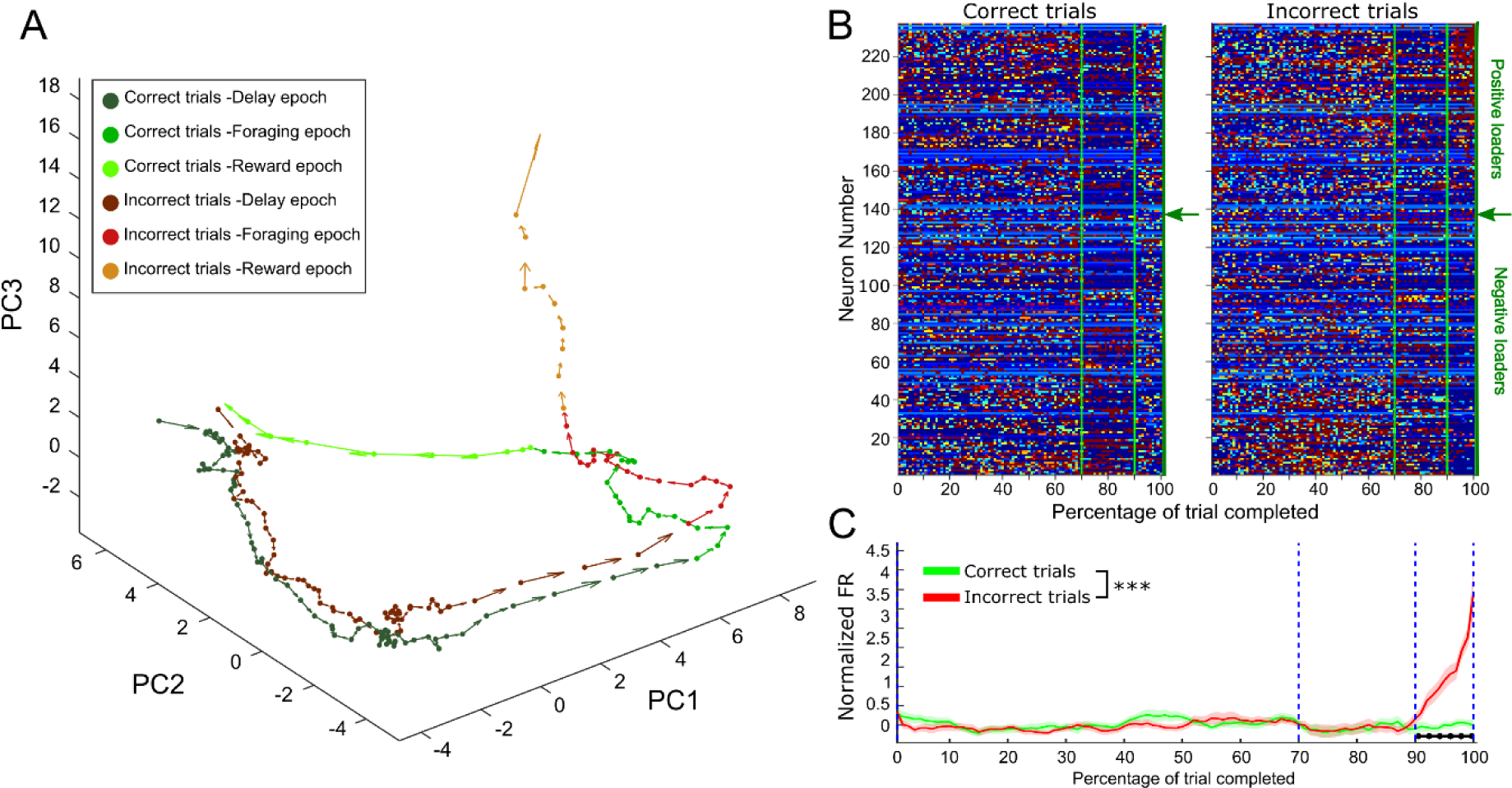
Divergence of the neural trajectory following an incorrect choice. A- Neural activity trajectories in the PC space for 237 pPys during correct and incorrect trials. Arrows indicate module and direction of trajectories’ speed. Different epochs of the task are color coded, transition between foraging and error epochs corresponds to a Correct dig for the correct trials and to an Error dig for the incorrect ones. PCA was performed on trial-normalized firing rates, the first 3 PC’s explained 38.5% of variance. B- Task-normalized firing rates for 237 pPys sorted according to their loadings on PC3 for correct (left panel) and incorrect (right panel) trials. Green vertical lines mark the End of the delay and the Dig event. Green arrows on the right side of each color-plot indicates the transition point between positive and negative loaders. C- Grand-average (±SEM) of task-normalized firing rate for the top 30% positive loaders on PC3 (30 pPys) on correct and incorrect trials. Firing rates in the two groups were significantly different (2-way ANOVA, interaction between kind of trial and time bin, F(99, 5800) = 1.59, p=2.0×10^−4^). Black horizontal markers indicate groups of time bins showing significant differences between correct and incorrect trials (FDR-corrected rank-sum, p<0.05).

## Discussion

The present experiment is the first to directly measure patterns of neural activity during the OST. We assessed activity in mPFC given its established role in the OST and other tests of working memory. The main findings of the study are: 1) Span lengths were bimodal and longer spans were associated with differences in neural activity of putative pyramidal neurons during the delay; 2) Sharp transitions in neural activity patterns emerge during the performance of the task that correspond to the onset of each behavioral epoch; 3) A transition was especially pronounced at the beginning of the foraging epoch where a group of putative interneurons were transiently and robustly active; 4) During foraging, neural activity patterns in putative pyramidal neurons were more robust during approach/digging of a novel than a familiar bowl; 5) A group of putative pyramidal neurons becomes active following an incorrect choice. Collectively, these data highlight the rich and evolving dynamics in mPFC that emerge throughout the performance of the OST. Therefore, the contribution of the mPFC to the OST is likely broad and diverse and not limited to the maintenance of a working memory.

### Neural activity at the termination of the delay correlates with span capacity

Our analyses of mPFC neural firing during the OST revealed complex patterns of neural activity that evolved throughout each of the epochs. Neural activity was increased in a subpopulation of putative pyramidal neurons during the delay, which then decreased sharply at the beginning of the foraging epoch (Figure 3, 5-B). Similar increases in mPFC delay activity that also predicted task performance have been reported in other spatial WM tasks (Myroshnychenko et al. 2017). In the current study, neural recordings were acquired in well-trained rats that likely anticipate the end of the delay, therefore, these changes in activity may reflect preparation for foraging. In addition, we observed that on high span trials, neural activity patterns at the end of the delay progressively became more similar to neural activity patterns observed during foraging (Figure 4-A, 5-B). This phenomenon may provide a smooth transition of the network to the foraging state and possibly facilitate the maintenance of information across the transition of the delay to the foraging epoch.

Putative interneurons were identified by their extracellular waveforms and most predominantly positively loaded on PC1 (Figure 2,3). Interneurons exhibited a rapid and pronounced increase in activity during the transition from the delay epoch to foraging epoch. This increase in activity was commensurate with increases in activity of a subpopulation of putative pyramidal neurons during this time as well. However, this was the only time during the task when transient increases in interneuron activity were detected, thus raising the question of what types of computations they facilitate during the transition from the delay to foraging epochs. During the foraging epoch, animals are required to sample different odors and refrain from digging in ones previously encountered. Effective execution of this part of the task requires that a representation of the previously visited bowls be held online in memory. This is extremely memory intensive and requires that between 4 and 17 odors be accessible in memory. Retention of these items in memory would be facilitated by a neural code that is flexible (e.g., it can be readily activated and inactivated) and highly dimensional (e.g., has high capacity). Maintaining a tight balance between excitation and inhibition in cortical networks has been suggested to facilitate an efficient yet high capacity coding scheme (Denève and Machens 2016). Therefore, we hypothesize that the concomitant increases in pyramidal and interneuron firing provide a network state that is capable of facilitating the transient maintenance of potentially large memory sets, such as those required to hold previously encountered odors in working memory. While there is precedent for this idea (Harvey et al. 2012; Lim and Goldman 2013, 2014), a more direct test of this hypothesis will provide critical clues as to how mPFC flexibly adapts to meet the computational needs of a given behavior.

### Neural activity in mPFC signals approach to novel, but not familiar, odors

The OST also has elements of a test of novelty detection whereby responding must be inhibited to familiar odors and then initiated (i.e., a dig) whenever a novel odor is detected. This pattern of responding requires maintenance of ‘familiarity’ for odors that have been experienced during the daily session and inhibition of digging when they are approached. We did not find evidence that a familiarity signal and/or inhibition of digging signal is maintained in mPFC on approaches to familiar odors. This was a surprising observation given reports of deficits in response inhibition following lesions of the mPFC (Miller and Cohen 2001; Fuster 2008; Chudasama 2011; Dalley et al. 2011). This suggests that errors in the OST following lesions of the mPFC may not be associated with computational processes required to inhibit behavior but rather incorrectly identifying an odor as novel.

Neural activity was strongly modulated during approach to a novel odor. This could be interpreted as a novelty signal which then triggers a dig in the correct bowl. The anterior cingulate cortex in humans/primates is proposed to code associations between rewards and actions, and in particular determine actions necessary to obtain rewards (Rushworth et al. 2011). Further, the mPFC in rodents may be involved in ‘working-with-memory’ during WM tasks, a function that optimizes behavioral responding during these tasks (Horst and Laubach 2012). As changes in neural activity were observed through approach, then digging, and retrieval of reward, they might also encode a more general signal that reflects the change in the behavioral requirements of the task during this epoch (e.g., stop foraging, dig, and retrieve reward).

Neural activity associated with approaches and initial digging on correct and incorrect trials did not differ. However, at the end of the digging, when the food pellet should be retrievable, robust differences in neural activity were observed. Upon receiving the food pellet on correct trials, neural activity patterns were qualitatively similar to those observed during the delay period. This is not surprising since the reward epoch signals the beginning of the delay. However, incorrect trials were uniquely characterized by a group of putative pyramidal neurons that increased firing when the animal would have been rewarded on a correct trial. It is possible that these neurons encode an “error” signal driven by the expectancy mismatch of expecting food and not receiving it. However, an incorrect dig, also signaled the end of the task for that day. As recordings were performed in well trained animals, it is likely the animals understood this, and this the signal may reflect environmental changes associated with the end of the task (e.g. being taken from the arena, etc.).

### Implications for theories of mPFC function during working memory and foraging

A number of neural activity patterns emerged throughout the OST. Robust transitions in the pattern of active neurons were observable across each behavioral epoch, which reflects a transiently stable population of (in)active neurons likely necessary to carry out the cognitive demands of each epoch. Similar phenomena have been observed in the mPFC of rodents engaged in foraging tasks (Lapish et al. 2008; Balaguer-Ballester et al. 2011) and operant tasks (Hyman et al. 2017) that require working memory. These “metastable” states are thought to provide an important mechanism to organize activity across populations of neurons to optimize information processing (Balaguer-Ballester et al. 2018). In the current study, the transitions between these states may have facilitated the updating of action plans required between each behavioral epoch.

The OST has been proposed as one of the few tasks suitable for measuring WM capacity in rodents and therefore provides an opportunity for identifying the brain mechanisms that underlie WM. Given the impairments in WM capacity seen in numerous brain disorders, use of this the task may provide an opportunity to model WM deficits in rodents and develop novel treatment approaches. Indeed, the OST shares some features in common with span tasks used to measure WM capacity in humans. However, differences between the OST and other WM capacity tasks used in humans and primates are notable. For example, a long delay period (at least in the context of WM) exists between the addition of each novel bowl in a given session and rats achieve odor spans much higher than the typical WM capacity limits in humans or primates. These differences have led some authors to question the specific nature of the cognitive function(s) measured by the task (April et al. 2013; Dudchenko et al. 2013; Branch et al. 2014; Davies et al. 2017). This study highlights the diverse and evolving patterns of neural activity observed in the mPFC of rodents performing the OST. Future research assessing the necessity of the observed neural activity states for span will increase understanding of the mPFC’s contribution to cognition, including those operations required for WM tasks with a significant capacity component.

## Materials and Methods

### Subjects

Seven adult male Long-Evans rats (220-250 g at arrival, Charles River, Quebec, Canada) were used. Rats were paired housed for one week in standard ventilated plastic cages on a 12 h light/dark cycle (lights on at 07:00). Food and water were available *ad libitum*. Rats were switched to individual cages, food restricted, and handled for approximately 5 min per day for 5 days before training commenced. Body weight was maintained at 85-90% of free feeding weight throughout the behavioral tasks. All experiments were performed according to the Canadian Council on Animal Care and were approved by the University of Saskatchewan Animal Research Ethics Board.

### Behavioral apparatus

Training and testing occurred on a 91.5 cm^2^ black corrugated plastic platform with 2.5 cm tall border. The platform was fastened to a metal frame with casters attached and stood 95 cm above the floor. It was surrounded by a beige curtain to block visual cues in the testing room. A Plexiglas box with a swinging door was placed in one corner of platform. Rats began each session in the box and were trained to go back to the box after obtaining reward for the delay period. The door was opened when trials started and closed when rats ran back into the box. Pieces of Velcro were equally spaced along the edge of the platform and used to fasten the sand filled bowls to the platform and stop the rats from spilling the sand. The bowls for a given trial were placed randomly on the pieces of Velcro. Odors were mixed in Premium Play Sand (Quikrete Cement and Concrete Products, Atlanta, GA) and then placed in white porcelain bowls (4.5 cm high, 9 cm in diameter) on the platform as needed for each trial. Sand (100 g) was scented by mixing 0.5 g of a single dried spice purchased from a local grocery store allspice, anise seed, basil, caraway, celery seed, cinnamon, cloves (0.1 g), cocoa, coffee, cumin, dill, fennel seed, garlic, ginger, lemon and herb, marjoram, mustard powder, nutmeg, onion powder, orange, oregano, paprika, sage, and thyme. The order of the odors used each day was selected randomly and rats were exposed to all odors many times before recordings began.

### Pre-surgery training

Dig training: Rats were trained to dig for a food reward (Kellogg’s Froot Loop) in a bowl filled with 100 g of unscented sand. Rats were placed opposite to a bowl on the platform for three separate phases. In the first phase, the food reward was positioned on top of the sand, in the second phase the food reward was incompletely buried, and in the third phase, the food reward was fully buried in the sand. Rats were trained until they would consistently dig for the food reward regardless of bowl position on the platform. This phase of training took 6–9 days to complete.

Delayed non-matching-to-sample (DNMS) task: In the sample phase (Trial 0), the rat was presented with a bowl of scented sand randomly positioned on the platform. Once the rat retrieved the Froot Loop, it was gently guided back to the box for the delay (40 s). In the choice phase of the trial (Trial 1), previously presented bowl was randomly re-positioned and a second bowl with a different odor was placed on the platform. A correct choice to obtain reward was recorded when rats dug into the bowl containing the novel odor. Rats moved on to the odor span task when they made 5 correct responses in 6 trials for 3 days.

Odor span task (OST): Trials of the OST (Figure 1-B) were run as described for DNMS task except that bowls with novel odors were added in subsequent trials until rats made an error (i.e., dug in any of the bowls except the novel one). The delay was maintained at 40 s. To prevent spatial cues from influencing performance, previous bowls were randomly positioned for each trial. Once rats achieved a span of 7 for two training days (8–16 days of training), the electrode array was implanted.

### Electrode implantation

Probes were custom built and consisted of a 4 × 8 matrix of tungsten wires (25 µm, California Fine Wire, Gover Beach, CA) in 35 Ga silica tubing (World Precision Instruments, Sarasota, FL). They were then attached via gold pins to an EIB-36-PTB board (Neuralynx, Bozeman, MT). Impedance to 200-600 kΩ measured at 1 kHz (NanoZ, White Matter LLC, Seattle, WA). Before surgery, rats were anesthetized with isoflurane, placed in a stereotaxic apparatus, and the dorsal surface of the skull was exposed. Four or 5 jeweler’s screws were threaded into the skull. Electrode arrays were then slowly lowered into medial prefrontal cortex (AP + 3.5–3.8 mm, ML ± 0.5 mm to the bregma, DV - 3.5 mm from the dorsal surface of the brain). A stainless-steel wire served as the ground and was soldered onto one of the skull screws dorsal to the cerebellum. Dental acrylic was used to secure the electrode array to the skull and screws. Rats were treated with Anafen immediately following the surgery and allowed to recover for 14 days from surgery before being retrained on the OST.

### Electrophysiological recordings

Rats were re-trained on the OST until their spans were higher than 4 for 2 days in row. Once this criterion was met, electrophysiological recordings were initiated during daily OST sessions. Rats were connected to a Digitalynx recording system controlled by Cheetah acquisition software (Neuralynx). Unit signals were recorded via a HS-36 unit gain headstage mounted on animal’s head by means of lightweight cabling that passed through a commutator (Neuralynx). Unit activity was amplified, sampled at 32 kHz, and bandpass filtered at 600–6,000 Hz. Local field potentials (LFPs) were sampled at 32 kHz and filtered at 0.1–9,000 Hz from each electrode. To verify the stability of recording, unit activity was recorded for about 15 min before and after the behavioral session. Behavior of the rats during the OST was monitored by a camera mounted to the ceiling with the experimental time superimposed on the video for offline analysis by a trained observer. Timestamps corresponding to trial start (when hind paws exited the Plexiglas box), delay start (re-entry to the Plexiglas box), familiar approach, novel approach, dig (forepaws contacting the sand), reward, and errors were recorded for each OST session.

### Histological verification of electrode positions

After the completion of all recording sessions, rats were deeply anesthetized with isoflurane, electrode positions were marked by electrolytic lesions (10 μA current for 10 s), and then the rats were perfused transcardially with physiological saline followed by 10% formalin. Brains were removed and post-fixed in a 10% formalin-10% sucrose solution. Brains were sectioned on a sliding microtome and infusion sites were determined using standard protocols with reference to a rat brain atlas (Paxinos and Watson 2006).

### Unit isolation and criteria

Spike sorting was performed offline with SpikeSort 3D, using a combination of KlustaKwik and manual procedures. Multiple parameters (spike height, trough, and energy) were used to visualize the clustered waveforms. Each cluster was then checked manually to ensure that the cluster boundaries were well separated, and waveform shapes were consistent with action potentials.

### Data analysis

Task normalized firing rates: For each correct trial, the three main epochs of the task (Delay, Foraging, and Reward epoch) were identified through the four behavioral timestamps: Delay start; Delay end; Correct dig; and End of trial (the latter corresponding to the Delay start marker of the following trial). A fixed percentage of trial completion was associated to each epoch by assigning a specific number of time bins (N) to it. Specifically, 70 time bins were assigned to the delay epochs, 20 to the foraging epochs, and 10 to reward epochs, for a total of 100 bins (see Figure 2-A2). The spike trains in each trial occurring between the beginning (T_j1_) and end time (T_j2_) of a specific epoch *j* were binned into N_j_ time intervals of duration t_j_:

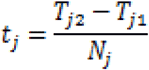

For a given neuron *i*, the firing rate in Hz along epoch *j* was obtained by dividing its corresponding bin counts for the time interval t_j_:

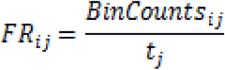

The single neuron’s firing rates across all epochs and trials were then z-score normalized. Finally, for each neuron, a task-normalized firing trace describing it’s mean activity during the execution of the task, was obtained by averaging across trials. The task-normalized firing rates on the error trials were obtained following a similar procedure, replacing the Correct Dig marker with the Error dig one, and setting the End of Trial marker at 10 s after the error dig. A separate analysis was performed using a fixed binning procedure to ensure that the results reported herein were not attributable to the task-normalized binning procedure (data not shown). Comparisons between the fixed and task normalized binning procedures lead to identical conclusions.

Waveform-based classification putative interneurons and pyramidal neurons: To separate putative interneurons from pyramidal neurons, single units were classified based on their average action potential (AP) waveforms using a clustering protocol proposed by (Ardid et al., 2015). For each of the 382 single units considered, action potentials were averaged and normalized in amplitude between −1 and 1. Each average waveform was interpolated with a cubical spline (from 32 original samples to 320 interpolated samples over 1ms time interval). Two features of the resultant waveform were then measured: the peak-to-trough duration and the time for repolarization (time, after the peak, to reach 25% of peak amplitude). Using principal component analysis (PCA), we integrated these two features into the first principal component (explaining 84% of total variance). The distribution of first components was tested for bimodality using a calibrated Hartigan’s dip test (Cheng and Hall 1998) (D(152) = 0.036, p=6.2×10^−3^). We fit the distribution with two Gaussian models and defined cutoffs to separate the two groups of narrow and broad waveforms (see Figure 2-B1). The two cutoffs were defined as the points at which the likelihood to belong to a group was 10 times larger than the likelihood to belong to the other one. Neurons with a principal component value smaller than the first cutoff (narrow waveforms) were classified as putative interneurons (pIn), while neurons with values larger than the second cutoff (broad waveforms) were classified as putative pyramidal cells (pPy). Neurons with a first component value falling between the two cutoffs were initially left unclassified. In several cases the AP waveform did not reach the repolarization threshold within the number of samples stored. Those cells were subsequently classified based only on their peak-to-trough duration which was compared to the peak-to-trough distributions for the classified waveforms (the peak-to-trough value had to exceed 5% confidence interval of the class distribution to be included in that class). Based on their average waveform (Figure 2-B2), the initially unclassified cells were subsequently merged with the pPy group. When looking at the mean firing rates of cells in each of the two classes (Figure 2-B3), we found that, as expected, the pIn population exhibited higher firing rates than the pPy one (Kolmogorov-Smirnov test, D(321,61)=0.30, p=1.1×10^−4^).

The task-normalized firing rates for pIns and pPys were compared through a 2-way analysis of variance (2-way ANOVA), where the interaction of cell class (pIn or pPY) and percentage of task completed (bins spacing from 1 to 100) was tested (interaction cell class x time, *F*(99,38000) = 3.02, *p* = 8.4×10^−22^). The firing rate between pIns and pPys at specific times were compared by re-binning the time-normalized data in 33 bins and differences in a given time bin were detected via FDR-corrected rank-sum tests (Figure 2-C). Identification of neural activity patterns via PCA: A principal component analysis was performed on the matrix of mean firing rates (*F*). Each column of the matrix *F* (100×382) contained the firing rate of a single neuron across the 100 time bins defining a trial of the task. We considered the first three principal components (PC) obtained, which, together, explained 56% of the total variance. The projection of the original data along the first three principal eigenvectors (Figure 3-A) identified the main neural patterns in our data. The task-normalized firing rates for the whole neural population were sorted according to their loadings on each of the first three PC’s (Figure 3-B). The loadings on each of the first three PC’s for pPys and pIns were compared using a Kolmogorov–Smirnov test (Figure 3-C).

Clustering of pyramidal neurons: In the PCA each cell receives a score (i.e., loading) for each PC, and therefore when classifying neurons based on a loading threshold it is possible for a neuron to be included in > 1 class. The goal of clustering pyramidal neurons based on their loadings was to group neurons into one class only for analyses. For this, PCA was applied to the task-normalized firing rates from the 321 identified pPy’s. Collectively, the first three PC’s explained 50.5% of the total variance and the respective loadings for each neuron were used as features in a k-means clustering algorithm. The optimal number of clusters was identified using the Akaike Measure of Information (Akaike 1974) adapted for k-means algorithm (Goutte et al. 2001) and defined as:

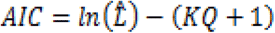

where the first term is the log-likelihood of the model (in our case, the specific clusters resulting from the k-means procedure), K is the number of clusters, and Q is the dimensionality (or number of features, 3 in our case). K-means algorithms can be considered as a form of Expectation Maximization algorithm for a Gaussian mixture model with equal weights and isotropic variances. The likelihood of our model was then estimated from the classification likelihood of each point *u*_*j*_, and it can be summarized in the equation:

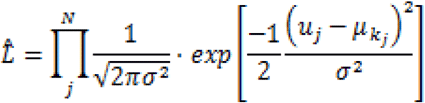

Where *N* is the total number of elements to classify, *σ^2^* is the average within-cluster variance calculated on all clusters, and *µ*_*kj*_ is the centroid of cluster *k* to which *u*_*j*_ is assigned. The AIC was calculated for values of k from 1 to 30 (Figure 5-A1), and a broken stick model was then used to select the number of clusters K=4 that optimally balanced information and compression.

Familiar vs novel odor approaches: Neural activities associated with approaches to familiar and novel odors were compared. Spike trains in a time interval of 4 s around each approach event (from −2 to 2 s) were binned in 40 intervals (0.1 s each). Events closer than 2 s to each other, to the end of delay marker, or to an error event were discarded. For each neuron, the firing rates obtained were normalized by the mean firing rate of the unit, and then averaged across all familiar approach events (mean familiar firing rate, fFR), and across all novel approach events (mean novel firing rate, nFR). Neurons with a median number of spikes around the approach events smaller than 2 or with less than 6 trials available for both types of approaches were discarded, leaving N=188 neurons available for the following analysis. Firing rates were smoothed using a moving average with a span of 5 bins. PCA was applied to the concatenated firing rate matrices (Nx80, where the first 40 columns contained the fFR’s and the last 40 columns contained the nFR’s). From the projections of firing rates along the first three PC’s we obtained the trajectories and speeds of the whole neural population around both familiar and novel approaches in the PC space (Figure 6-A). For the following analysis we only included time bins up to 0.3 s after an approach. This was done to avoid contamination in the activity coming from either the dig or the reward; a correct dig happened before 0.3 s from a novel approach only in 1.8% of the trials considered (15 out 823). By concatenating the matrices vertically (2Nx23), a second PCA provided 2 loading coefficient sets for each neuron (one for familiar and one for novel approaches). Firing rates for the positive and negative loaders on each PC for the two approaches were grouped and averaged (Figure 6-B). Distribution of absolute loadings on the first 3 PC’s for the two approaches were compared by means of the Kolmogorov-Smirnov test (Figure 6-C). Incorrect choice trials: In 67 out of the 77 sessions considered for analysis an incorrect choice trial was also recorded. Incorrect choices occurred when the animal dug into a non-novel odor bowl and it resulted in the session ending. Firing activity during a single incorrect trial was available for each of the 237 pyramidal neurons recorded from these sessions. Task-normalized firing rates for the incorrect trials were obtained between the four behavioral timestamps: Delay start; Delay end; Error dig; and End of trial and arranged in a matrix of size 237×100, where each row corresponded to the activity of a single neuron. For the same neurons, a random correct trial was selected for comparison, and the related task-normalized firing rates were arranged in a second matrix of size 237×100. The two matrices were concatenated row wise and PCA was applied to the resulting 237×200 matrix. PCA space trajectories and speeds of the neural population on correct and incorrect trials were then obtained from the projections of firing rate matrices along the first three PC’s (Figure 7-A), which together explained 38.5% of the variance. Note that a single correct trial was selected for this procedure to keep the signal’s noise comparable in the two matrices. As a control for possible effects due to the trial’s order, we also tried to select the random correct trial among the last 5 correct trials. PCA space trajectories obtained in this case were similar to those obtained with the unconstrained selection of the random correct trial. Task-normalized firing rates for the 237 pPys were sorted according to their loadings on PC3 (Figure 7-B). Firing rates for the top 30% positive loaders on PC3 were compared in the two conditions (correct and incorrect trials) through a 2-way analysis of variance. The firing rates at specific times were compared via FDR-corrected rank-sum tests, as described for Figure 2C.

## Acknowledgements

We thank Jillian K. Catton and Alex Senger for technical assistance with this project and Jeremy K. Seamans and James M. Hyman for comments on the manuscript.

